# Connectivity between the hippocampus and default mode network during the relief – but not elicitation – of curiosity supports curiosity-enhanced memory enhancements

**DOI:** 10.1101/2021.07.26.453739

**Authors:** Charlotte Murphy, Charan Ranganath, Matthias J. Gruber

**Affiliations:** Cardiff University Brain Research Imaging Centre (CUBRIC), School of Psychology, Cardiff University, Wales CF24 4HQ, United Kingdom; Department of Psychology, University of California at Davis, Davis, California 95616, USA; Center for Neuroscience, University of California at Davis, Davis, California 95616, USA

## Abstract

Consistent with the idea that curiosity enhances information seeking, it has been shown that activity within both the dopaminergic circuit and hippocampus supports curiosity-enhanced learning. However, the role of whole-brain mechanisms involved in cognitive control (fronto-parietal network; FPN) and memory integration (default mode network; DMN) that might underpin curiosity states and their effects on memory remain elusive. We hypothesised that the FPN and DMN should distinguish between high- and low-curiosity conditions and be recruited more heavily for later remembered information associated with high-curiosity. Here, we used functional magnetic resonance imaging whilst participants completed a trivia paradigm, in which we presented trivia questions associated with high- and low-curiosity, followed by the associated answer. After a short delay, we tested memory for trivia answers. We adopted a network-based parcellation of the brain into subnetworks of the FPN and DMN to examine how neural activity within, and functional connectivity between, each subnetwork predicts curiosity-enhanced memory. Across elicitation and relief of curiosity, we found focal recruitment of FPNA and widespread recruitment of DMN subnetworks in support of curiosity and curiosity-enhanced memory. Most importantly, during the elicitation of curiosity, functional subcortical connectivity and across cortical networks, but not subcortical-cortical coupling, correlated with curiosity-enhanced memory. However, during the relief of curiosity, coupling between subcortical regions and DMNA emerged in support of curiosity-enhanced memory. Taken together, our results provide the first evidence about how neuromodulatory mechanisms via the hippocampal-dopaminergic circuit trigger states of curiosity and thereby communicate to higher-order cortical regions to facilitate curiosity-enhanced memory.

**Significant statement:** Does neural activity within, and functional connectivity between, the dopaminergic-hippocampal network, fronto-parietal network (FPN), and default mode network (DMN) underpin curiosity states and their effects on memory? Here, we show how the dopaminergic system together with the hippocampus interact specifically with subnetwork DMNA potentially reflecting how subcortical regions support the enhancement of memory intergration of semantic information associated with curiosity. As DMNA (the core DMN subnetwork) was also functionally coupled with the whole DMN network and the semantic control network (FPNA), these findings provide a plausible neuromodulatory mechanism through which hippocampal-dopaminergic input triggers curiosity and then communicates to higher-order brain regions via DMNA to facilitate curiosity-enhanced memory.

## Introduction

A growing body of research that addresses the relationship between curiosity and learning uses a trivia paradigm in which participants are tested on memory for answers to trivia questions that elicit different levels of curiosity (e.g., Kang et al., 2009). These studies demonstrate that memory of trivia answers is higher for questions that elicited high levels of curiosity (coined from here-on out as curiosity-enhanced memory) (Fastrich et al., 2018; Galli et al., 2018; Marvin & Shohamy, 2016; McGillivray et al., 2015; Mullaney et al., 2014; Murayama et al., 2010; Stare et al., 2018; Wade & Kidd, 2019). At the meso-scale, research suggests that curiosity states are related to modulations in activity in the hippocampus-dopaminergic circuit and these modulations impact memory encoding for curiosity target information (Murayama, 2019; Oosterwijk et al., 2020; Sharot & Sunstein, 2020). For example, it has been shown that during the presentation of stimuli that elicit curiosity (i.e., trivia question presentation), activity in the dopaminergic midbrain regions (e.g., substantia nigra/ventral tegmental area: SN/VTA) and the hippocampus predicts curiosity-related memory enhancements for curiosity target information (Gruber et al., 2014).

Further functional magnetic resonance imaging (fMRI) studies have demonstrated stronger activation of the inferior parietal lobule during the elicitation of curiosity (van Lieshout et al., 2018). Convergently, curiosity-enhanced memory of trivia answers was associated with increased activity in inferior parietal lobule, middle temporal gyrus and prefrontal cortices (Duan et al., 2020). In addition, perceptual curiosity seems to engage frontal and parietal areas implicated in attention and cognitive control (Jepma et al., 2012). These regions form the frontoparietal network (FPN), which is thought to be critical for our ability to coordinate behaviour in a flexible goal-driven manner (Marek & Dosenbach, 2018) and has been suggested as a possible candidate for mediating curiosity states based on its connection to reward-related midbrain regions (Gottlieb, Lopez & Baranes, 2013; Gottlieb & Oudeyer, 2018). Additionally, curiosity-enhanced activity encompassing angular gyrus and medial prefrontal cortex was also found in Duan et al. (2020) during the relief of curiosity (i.e., trivia answer presentation). Collectively, these brain regions form part of the default mode network (DMN) - a key cortical network involved in memory. Multiple lines of evidence suggest that BOLD signal fluctuations in the DMN are functionally tightly connected to the SN/VTA and hippocampus (Bär et al., 2016; Ranganath & Ritchey, 2012; Ritchey, Libby & Ranganath, 2015). Recent attempts have been made to investigate the relationship between trait curiosity and functional connectivity across the FPN and DMN (Li et al., 2019). However, direct investigation of the role of both the FPN and DMN, and their connection with the SN/VTA and the hippocampus, in supporting states of curiosity and related memory enhancements is lacking.

The current study combines a trivia question task (Figure 1) in conjunction with functional magnetic resonance imaging (fMRI) to address several outstanding questions: (i) Do subnetworks of the FPN and DMN support curiosity and curiosity-enhanced memory? (ii) Do the SN/VTA and hippocampus show functional coupling with such cortical subnetworks to support curiosity-enhanced memory? (iii) Are these subnetworks recruited differentially during curiosity elicitation and relief? We predicted that FPN and DMN should distinguish when participants are in a high-compared to a low-curiosity state and be recruited more heavily for high-curiosity trials that are subsequently remembered. Furthermore, we used task-based functional connectivity analysis to ask whether and how the hippocampus and SN/VTA – previously shown to support curiosity-enhanced memory – show functional coupling with cortical brain networks (i.e., subnetworks of the FPN and DMN) in order to support states of curiosity and their effects on memory. We predicted that the hippocampus and SN/VTA will show greater functional coupling to subnetworks within the FPN and DMN during states of high curiosity when items are subsequently remembered. We also addressed the question whether hippocampus-SN/VTA coupling with FPN/DMN will be most prominent during elicitation or relief of curiosity.

**Figure 1.**
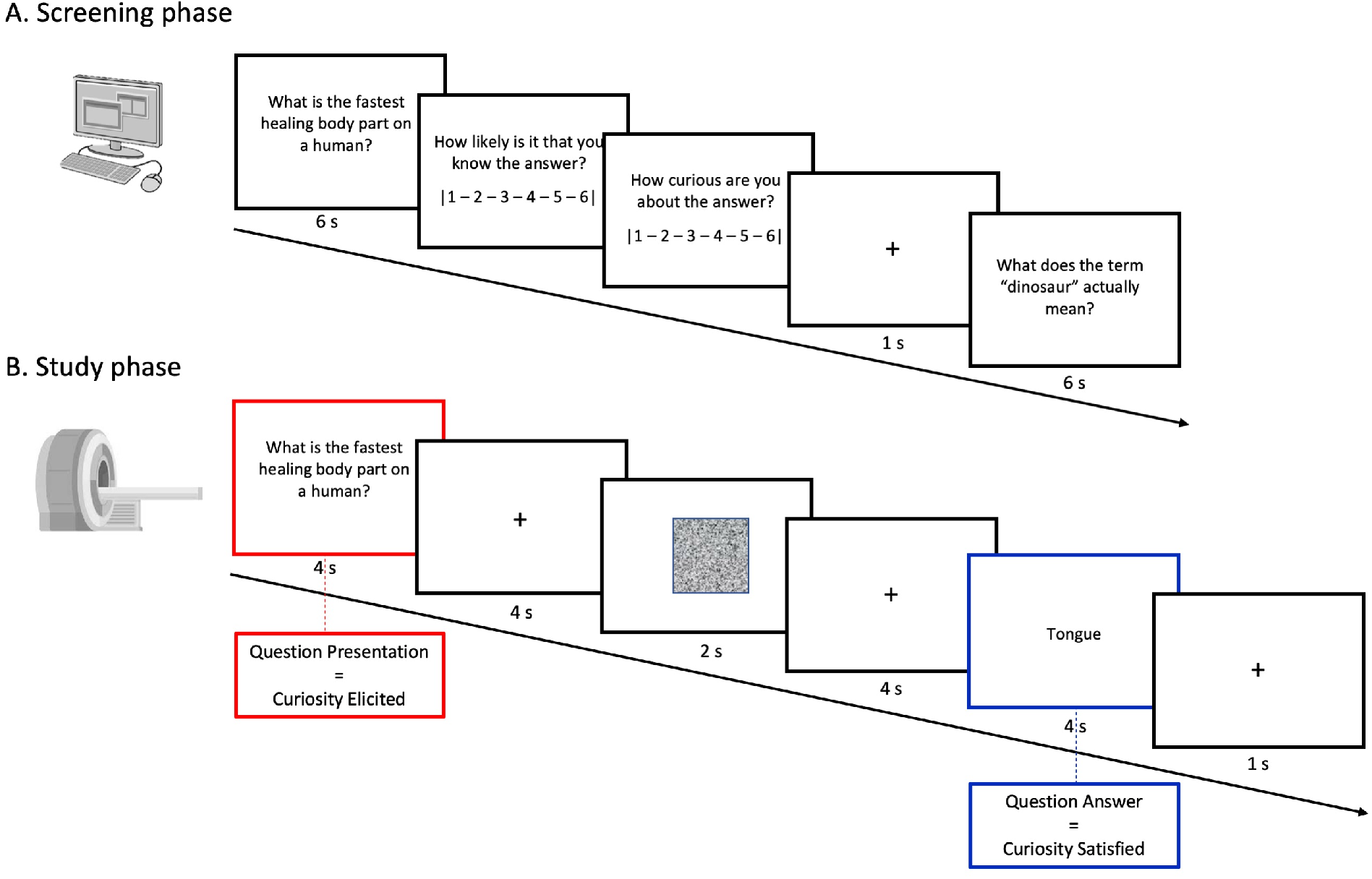
Experimental design of screening and study phases of the trivia paradigm. (A) Screening phase (performed on a computer): for each trial, participants rated how likely they knew the answer to a trivia question and how curious they were to learn the answer. 112 questions associated with high and low curiosity, for which participants did not know the answer, were used for the next phase. Answers were not presented in this phase. (B) Study phase (performed in the MRI scanner): For each trial, a selected trivia question was presented, and the participant anticipated the presentation of the answer. During this anticipation period, participants were required to make an incidental judgment to a face (shown as colour image in the experiment). As the purpose of the current analysis was to look at differences between the elicitation and satisfaction of curiosity this period was not analysed and so denoted by a grey box. Following the study phase, participants completed memory tests (not shown) on both the trivia answers and the faces that were studied in the scanner.

## Material and methods

### Participants

Twenty-four healthy young adults took part in the experiment. Six participants were excluded from the analyses due to the following reasons: one participant due to excessive movement artifacts in the fMRI data, two participants due to failure to comply with the instructions, and three participants because of insufficient number of trials for one participant. Results are based on the remaining eighteen participants. Participants’ mean age was 22.4 years (range: 18-31). Seventeen participants were right-handed and one left-handed. They were compensated with $50 for their total time in the laboratory. All participants had normal or corrected-to-normal vision and were native English speakers. The UC Davis Institutional Review Board approved both experiments. This data has been used in a previous publication (Gruber et al., 2014).

### Experimental Design

#### Overview

This is a reanalysis of published data (Gruber et al., 2014), where participants underwent a four-stage paradigm with (1) a screening phase, (2) a study phase, (3) a surprise recognition test phase for incidentally presented faces, and (4) a surprise recall test for trivia answers presented during the study phase. The delay between the study phase and the memory test was on average 20 minutes.

### Stimuli

#### Trivia questions and answers

We generated a pool of 375 trivia questions along with their corresponding answers from online trivia websites. Questions corresponded to the following trivia categories: history/geography, movies/TV, music, nature, science, space, sports, food, and other miscellaneous facts. The pool only included trivia questions for which the answers were likely to be unknown to the majority of participants because participants should not have prior knowledge but learn the answers during the study phase. On average (min-max), trivia questions contained 11 (4-20) words and trivia answers 2 (1-8) words. The selected 112 trivia items for the study phase differed across participants because trivia questions were randomly drawn from the trivia pool in the screening phase and the allocation of trivia questions to high- or low-curiosity conditions depended on participants’ ratings during the screening phase.

### Design

Throughout all phases of the experiment, stimuli were presented on a grey background and in the centre of the computer screen. The Psychophysics Toolbox (http://psychtoolbox.org) was used for the presentation of all stimuli. In the screening phase, a trivia question was presented for 6 s followed by two consecutively presented rating scales that were self-paced (see Figure 1A). After a response was given for the second rating, an inter-trial fixation cross was presented with a duration of 1 s. In the study phase (Figure 1B), a trial started with a trivia question that was presented for 4 s and ended with a 1 s long presentation of the trivia answer or the letter string ‘xxxxx’ on catch trials. During the 14 s long anticipation phase (i.e. from the onset of the trivia question to the onset of the trivia answer), a cross hair was presented after the presentation of the trivia question and the cross hair was replaced by a face from 8 to 10 s after the onset of the trivia question. A cross hair was also presented during the inter-trial interval, which was temporally jittered with an average of 4 s within a scanning run. In the recall test phase for trivia answers, a Microsoft Excel spreadsheet was presented that included one column with a random order of all 112 trivia questions presented during the study phase and participants were instructed to fill in the answers in the column right next to the question.

### Procedure

#### Screening Phase

Because the level of curiosity elicited by different trivia questions varies between participants, we used participants’ ratings to sort trivia questions into participant-specific high- and low-curiosity categories (56 questions each). Trivia questions were randomly selected from a pool of 375 trivia questions and were consecutively presented. After the presentation of a trivia question, participants had to give two self-paced ratings on 6-point scales (see Figure 1A). First, they had to rate how confident they were that they knew the answer to a trivia question (extremes of scale: 1 = “I am confident that I do not know the answer” and 6 = “I am confident that I know the answer”). Second, participants rated their level of curiosity about the answer to a trivia question (extremes of scale: 1 = “I am not interested at all in the answer” and 6 = “I am very much interested in the answer”). If participants did not indicate that they knew the answer to a trivia question (i.e., they did not give a 6-point response on the answer confidence rating), trivia questions with response points 1–3 on the curiosity rating were allocated to the low-curiosity condition and response points 4–6 to the high-curiosity condition. The screening phase lasted until 56 trivia questions were allocated for each curiosity condition. On average (min-max), participants gave a high-curiosity rating on 85 (range, 56–173) and a low-curiosity rating on 58 (range, 56–68) trivia questions.

#### Study Phase

In the subsequent study phase that took place in an MRI scanner, the selected 112 trivia questions were presented along with the associated answers (see Figure 1B). A trial started with the presentation of a trivia question, followed by an anticipation period that preceded the presentation of the associated trivia answer. Six of the 56 trials (∼10%) in each condition were catch trials to ensure participants’ attention throughout the scanning session. In these trials, the letter string “xxxxx” was presented instead of the trivia answer. During the anticipation period, a cross-hair was presented that was replaced by an image of an emotionally neutral face (incidental item) during the middle of the anticipation period. During the presentation of the face, participants had to give a yes/ no response (on an MRI-compatible response box) as to whether this particular person would be knowledgeable about the trivia topic and could help them figure out the answer. The study phase was divided into four scanning runs (9 min each).

#### Recall Test for Trivia Answers

Approximately 20 min after the end of the study phase inside the MRI scanner, participants took part in two surprise memory tests outside of MRI scanner. First, a surprise recognition memory test for incidental faces was administered (note that memory for faces and the associated neural mechanisms are reported in our previous publication (Gruber et al., 2014) but were not analysed for the current manuscript). After the recognition memory test for faces, participants were given a list with all trivia questions from the study phase in random order. Participants were encouraged to take approximately 20 min to write down the correct answers without guessing any answers.

### fMRI acquisition

We used a 3T Siemens Skyra scanner with a 32-channel phased array head coil to acquire anatomical and functional MRI images. A multiband Echo-Planar Imaging sequence was used to acquire whole brain T2∗-weighted images (repetition time = 1.22 s, echo time = 24 ms; 38 slices per volume; multiband factor = 2; voxel size = 3 mm isotropic) with 441 volumes for each of the four scanning runs. In addition, a T1-weighted MP-RAGE with whole brain coverage was acquired. Inside the head coil, the participant’s head was padded to restrict excessive motion. Stimuli were displayed on a mirror attached to the head coil above the participant’s eyes. During the scanning, the participant’s eyes were monitored by the experimenter via an eye tracker to ensure that the participant attended to all stimuli.

### fMRI analysis

#### Software

Functional and structural data were pre-processed and analysed using FMRIB’s Software Library (FSL version 6.0, http://fsl.fmrib.ox.ac.uk/fsl/fslwiki/FEAT/).

#### ROI definition

17 whole-brain networks were defined based on a well-established 17-network parcellation (Yeo, Krienen et al., 2011). Guided by previous findings on curiosity-enhanced brain activation, we then selected all networks that encompassed the frontoparietal control network (FPN) and default mode network (DMN) which resulted in 6 regions-of-interest (ROIs) (shown in Figure 2). Two ROIs that encompassed the FPN [the Yeo et al. 17-network parcellation actually divides the unified FPN into three separate subnetworks; however, one subnetwork is only composed of two regions (the posterior cingulate and precuneus) and does not include a frontal component. As such, it is not a frontoparietal system per se, and is not examined here] and four that encompassed the DMN. These have been given the following names: FPNA = frontoparietal control network A, FPNB = frontoparietal control network B, DMNA = default mode network A, DMNB = default mode network B, DMNC = default mode network C and DMND = default mode network D (see Figure 2). For the functional connectivity analyses two subcortical ROIs were taken from a previous publication of this data. The hippocampus ROI was taken from (Gruber et al., 2016) and the dopaminergic ROI encompassing the substantia nigra / ventral tegmental area complex (SN/VTA) taken from Murty and Adcock (2014) (see Figure 4). This allowed us to determine cortical-subcortical connections. Specifically, it permitted the investigation of how subcortical regions (previously shown to be pivotal in capturing the elicitation of a curiosity state and subsequent memory effect; Gruber et al., 2014) communicate with cortical networks. These subcortical ROIs reflect bilateral hippocampus and bilateral dopaminergic SN/VTA.

**Figure 2.**
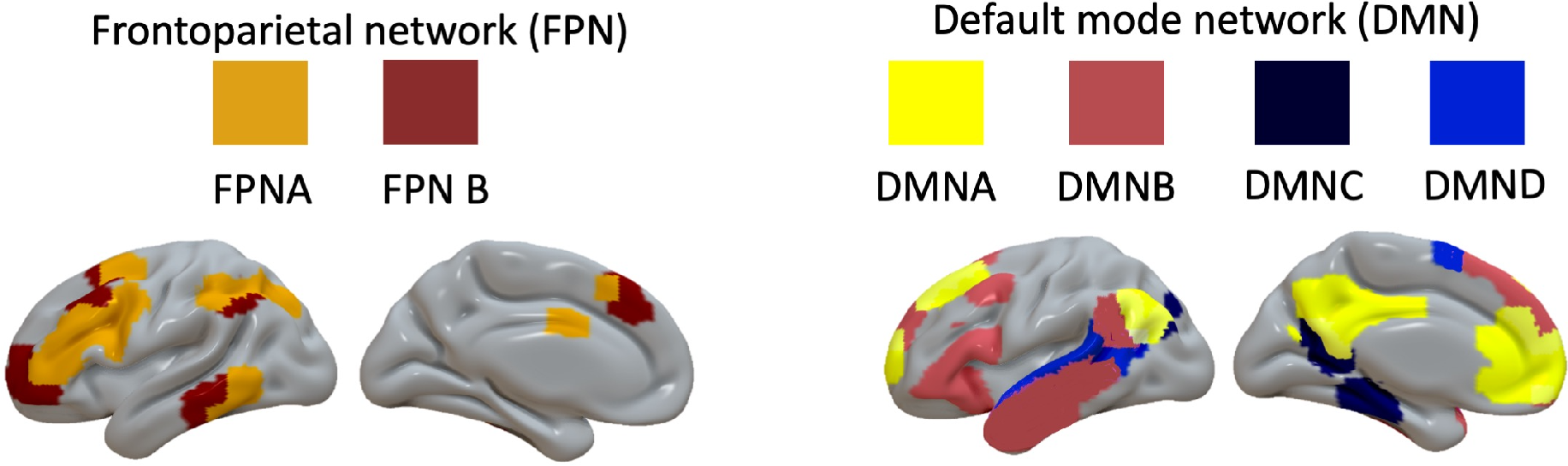
Regions of interest spanning the FPN and DMN. Using the 17-network parcellation by Yeo et al. (2011), we selected six ROIs that encompass the frontoparietal network (FPN) and default mode network (DMN).

#### Pre-processing

All imaging data were pre-processed using a standard pipeline and analysed via FMRIB Software Library (FSL Version 6.0). Images were skull-stripped using a brain extraction tool (BET; Smith, 2002). The first four volumes of each scan were removed to minimize the effects of magnetic saturation, and slice-timing correction with Fourier space time-series phase-shifting was applied. Motion correction (MCFLIRT (Jenkinson et al., 2002),) was followed by temporal high-pass filtering (cut-off = 0.01 Hz) and spatial smoothing (Gaussian full width half maximum 6 mm). Individual participant data was registered to their high-resolution T1-anatomical image, and then into a standard spare (Montreal Neurological Institute); this process included tri-linear interpolation of voxel sizes to 2 × 2 × 2 mm.

### Statistical analysis

#### Network ROI - Univariate analysis

General linear models (GLMs) were estimated by modelling BOLD signal changes using a stick function (0 s duration) to model the onset of the particular events. We convolved these stick functions with a double-gamma hemodynamic response function and included the 6 motion covariates to account for motion-related noise in the data (i.e., three rigid-body translation and three rigid-body rotation parameters). Catch trials were modelled separately for all event onsets and were not included in any analyses. 12 regressors were modelled in total; 3 x timing of presentation (trivia questions, faces, and trivia answers); 2 x curiosity level (high, low); 2 x memory performance for trivia answers (remembered, forgotten). Each run was modelled separately, resulting in four different models per participant. A fixed effect design (FLAME, http://www.fmrib.ox.ac.uk/fsl) was then conducted to average the four runs, within each individual.

Statistical analysis was performed using FSL and SPSS. Group-level analysis of multiple conditions was performed using repeated-measures ANOVA, with partial eta squared (η_p_^2^) reported as the effect size. The within-subject factors corresponding to each test are provided in Results. For each participant, activity was extracted from each of the 6 binarised ROIs (using FSLs Featquery command) and then entered into a group-level 2 x 2 x 2 repeated-measures ANOVA for each ROI with the following factors: Curiosity (2 levels; high, low) x Memory (2 levels; remembered, forgotten) x Timepoint (2 levels; onset of trivia question, onset of trivia answer). Although the face onset timepoint was modelled, the analyses of interest were the onsets of the trivia questions and answers (i.e., when curiosity was elicited and relieved) and therefore face onset was not included in the repeated-measures ANOVAs.

#### Functional connectivity analyses

As prior research has indicated that hippocampal-dopaminergic regions support curiosity-enhanced memory during elicitation, and the current univariate analysis revealed that default mode network supports curiosity-enhanced memory during satisfaction we next ran a functional connectivity analysis between our SN/VTA and hippocampus ROI with the 5 network-based ROIs that showed a curiosity-enhanced memory effect. For each of our five network ROIs we further parcellated these down into key regions by masking each subnetwork by binarized lateral and medial lobe masks (lateral frontal, medial frontal, lateral temporal, medial temporal, lateral parietal and medial parietal). The resulting ROIs were then given an anatomical label that referenced the central voxel of that region. This resulted in the following ROIs for each subnetwork: FPNA (SPL = superior parietal lobule, pITG = posterior inferior temporal gyrus, lPFC = lateral prefrontal cortex), DMNA (mPFC = medial prefrontal cortex, PCC = posterior cingulate cortex, dAG = dorsal angular gyrus), DMNB (FP = frontal pole, ATL = lateral temporal lobe extending into anterior temporal lobe, vAG = ventral angular gyrus), DMNC (LOC = superior lateral occipital cortex, Prec = precuneous, PHG = parahippocampal gyrus) and DMND (SMG = supramarginal gyrus, STG = superior temporal gyrus). The mean time-series was extracted from each ROI separately for each of our 8 conditions [2 x timepoints (question, answer), 2 x curiosity (high, low) and 2 x memory (remembered, forgotten)].

To determine which functional connections were associated with curiosity-enhanced memory Fisher-transformed bivariate regression coefficients (beta values) between two ROI BOLD time-series were used to identify significant increases or decreases in functional connectivity between the seed-to-target. Specifically, the times series for remembered – forgotten were computed for High and Low curiosity respectively at each time point, yielding four average time series per participant: (1) High Curiosity Question Presentation (remembered – forgotten), (2) High Curiosity Answer Presentation (remembered – forgotten), (3) Low Curiosity Question presentation (remembered – forgotten) and (4) Low Curiosity Answer Presentation (remembered – forgotten). Finally, as we were interested in the interaction between curiosity and memory (i.e., the functional coupling that predicts curiosity-enhanced memory) the following calculation was performed for each timepoint: High Curiosity (remembered – forgotten) – Low Curiosity (remembered – forgotten). Pearson correlations were calculated between the signal time-course of each participant in the first ROI, and the time course in the second ROI. This yielded a matrix of values per ROI pair, describing the fluctuation of coactivations among each pair of tested regions for each condition. This was done first at the whole-network level (Figure 4) and then parcellated into the key regions that make up each network (Figure 5). All *p*-values were adjusted to control for false-discovery rate (FDR) using the Benjamini Hochberg correction in R (Benjamin & Hochberg, 1995).

#### Code and data accessibility

All custom code, anonymised data and ROIs are publicly available on the Open Science Framework: https://osf.io/m4nu9/

## Results

### Behavioural findings show curiosity-enhanced memory

For consistency with the fMRI results, behavioural results are reported for the 18 participants that were used for the fMRI analyses. Consistent with findings in our previous publication using the same dataset (Gruber et al., 2014), participants recalled significantly more answers to high-curiosity questions compared to low-curiosity questions (70.6% *SE* = ±2.60 versus 54.1% *SE* = ±3.04; *t*(17) = 5.64, *p* < 0.001).

### FMRI results

#### What are the cortical networks that support curiosity?

To first determine which networks are more involved in high- compared to low-curiosity conditions, we examined the univariate BOLD activity during high- vs. low-curiosity trials within each of our six ROIs (see Figure 2). The results of our repeated-measure ANOVAs revealed a significant main effect of curiosity in three of our ROIs: FPNA, DMNB and DMNC (Table 1 – middle column).

**Table 1.** Results from repeated-measures ANOVAs investigating effects of Curiosity and Timepoint on BOLD activity.

| ROI | Network | Main Effect |  |  |  |  |  | 2-way interaction |  |  |
| --- | --- | --- | --- | --- | --- | --- | --- | --- | --- | --- |
|  |  | Curiosity |  |  | Timepoint |  |  | Curiosity x Timepoint |  |  |
| | | *F* | *p* | $\eta_p^2$ | *F* | *p* | $\eta_p^2$ | *F* | *p* | $\eta_p^2$ |
| 1 | FPN A | **12.20** | **.004 \*** | **.47** | 3.62 | .078 | .21 | 1.92 | .668 | .01 |
| 2 | FPN B | 2.66 | .125 | .16 | 0.01 | .915 | .00 | 0.48 | .498 | .03 |
| 3 | DMN A | 2.65 | .126 | .16 | 0.57 | .465 | .04 | 0.36 | .561 | .03 |
| 4 | DMN B | **5.84** | **.030 \*** | **.29** | **7.14** | **.018\*** | **.34** | 0.32 | .579 | .02 |
| 5 | DMN C | **8.10** | **.013 \*** | **.37** | **7.10** | **.018\*** | **.37** | 0.55 | .469 | .04 |
| 6 | DMN D | 3.40 | .087 | .20 | 0.02 | .900 | .00 | 0.59 | .456 | .04 |

*Note. Left-hand column: Main effect of curiosity for each ROI. Middle column: Main effect of timepoint for each ROI. Right-hand column: Two-way interaction between curiosity (high, low) x timepoint (question, answer) for each ROI. \* denote a significant interaction or main effect at $p < .05$ . $\eta_p^2$ = partial eta squared effect size.*

Next, to determine whether our ROIs support curiosity differentially during the elicitation (i.e., question presentation) and relief (i.e., answer presentation) of curiosity, we examined the interaction between Curiosity (high, low) x Timepoint (question, answer). The results revealed no significant interaction in any of our ROIs (Table 1 – left column), suggesting that these networks are recruited in comparable ways to support curiosity during the elicitation and relief of curiosity.

#### Which cortical networks support curiosity-enhanced memory for trivia answers?

To determine whether any of our network ROIs support curiosity-enhanced memory (i.e., higher activity of remembered vs. forgotten answers during high- compared to low-curiosity conditions), we examined the interaction between Curiosity (high, low) x Memory (remembered, forgotten). The results revealed that there was a significant two-way interaction between Curiosity x Memory in five ROIs: FPNA and all four DMN ROIs (Table 2 – middle column; Figure 3). These findings indicate that activation for subsequently remembered trivia answers compared to forgotten trials was significantly enhanced for high-curiosity trials in FPNA and all default subnetworks. Taken together, this indicates that curiosity-enhanced memory may be supported by focal recruitment of FPNA and widespread recruitment of default regions.

**Table 2.** Results from repeated-measures ANOVAs investigating effects of Curiosity, Memory, and Timepoint on BOLD activity.

|  | Network | 2-way Interaction |  |  |  |  |  | 3-way Interaction |  |  |
| --- | --- | --- | --- | --- | --- | --- | --- | --- | --- | --- |
|  |  | Curiosity x Memory |  |  | Curiosity x Timepoint |  |  | Curiosity x Memory x Timepoint |  |  |
| | | *F* | *p* | $\eta_p^2$ | *F* | *p* | $\eta_p^2$ | *F* | *p* | $\eta_p^2$ |
| 1 | FPN A | **5.06** | **.041 \*** | **.27** | 0.19 | .101 | .18 | 0.01 | .960 | .00 |
| 2 | FPN B | 1.76 | .205 | .11 | 0.48 | .498 | .03 | 0.01 | .936 | .00 |
| 3 | DMN A | **5.90** | **.029 \*** | **.30** | 0.36 | .561 | .03 | 0.45 | .514 | .03 |
| 4 | DMN B | **9.18** | **.009 \*** | **.40** | 0.32 | .579 | .02 | 0.99 | .336 | .07 |
| 5 | DMN C | **10.68** | **.006 \*** | **.43** | 0.55 | .469 | .04 | 0.03 | .865 | .00 |
| 6 | DMN D | **6.59** | **.022 \*** | **.32** | 0.59 | .456 | .04 | 0.24 | .630 | .02 |

*Note. Left-hand column: Main effect of curiosity x memory for each ROI. Middle column: Main effect of timepoint for each ROI. Right-hand column: Three-way interaction between curiosity (high, low) x memory (remembered, forgotten) x timepoint (question presentation, answer presentation) for each ROI.*
*\* denote a significant interaction or main effect at $p < .05$ . $\eta_p^2$ = partial eta squared effect size.*

**Figure 3.**
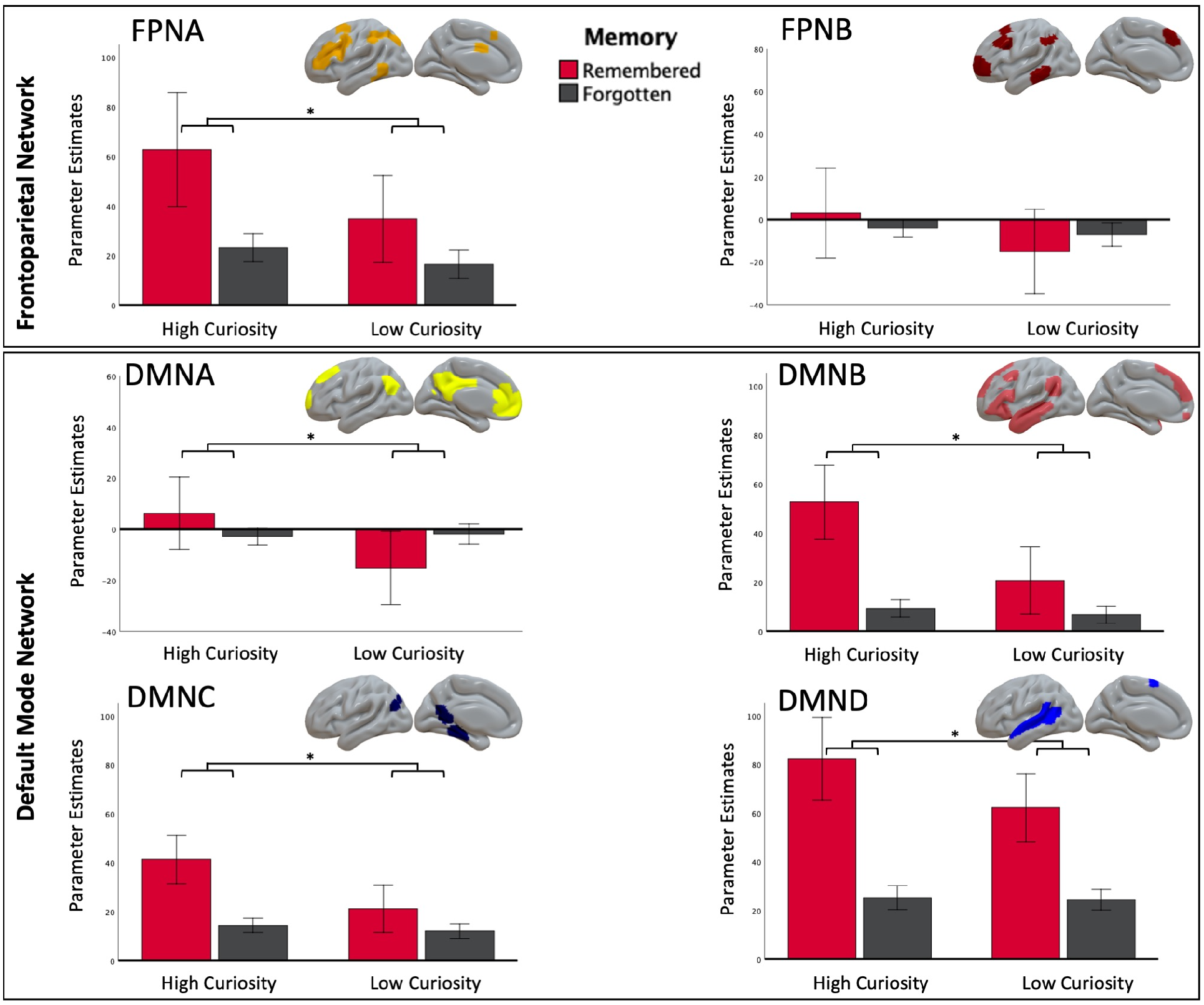
During both curiosity elicitation and relief, curiosity-enhanced memory for trivia answers is supported by focal recruitment of FPNA and widespread recruitment across all DMN subnetworks. BOLD activity for remembered (red bars) and forgotten (grey bars) trivia answers are presented on the x-axis for high and low-curiosity conditions for each ROI independent of timepoint (question, answer). Visualisation of each ROI network taken from the 17-network parcellation by Yeo et al. (2011) reveal five ROIs that show enhanced activity (FPNA, DMNA, DMNB, DMNC and DMND) during high-compared to low-curiosity states for successfully recalled trivia answers. Standard error of the mean represented by whisper bars.

Finally, to determine whether subnetworks of FPN and DMN contribute differentially to curiosity-enhanced memory during the elicitation and relief of curiosity, we investigated the three-way interaction between curiosity x memory x timepoint. The results from this repeated-measures ANOVA revealed that no ROIs showed a significant interaction between curiosity, memory, and timepoint (Table 2; left hand column). In line with our findings on curiosity-enhanced activity, our results suggest that the subnetworks supporting curiosity-enhanced memory (FPNA, DMNA, DMNB, DMNC and DMND) are recruited comparably during the elicitation and relief of curiosity.

#### How does functional connectivity between subcortical (i.e., SN/VTA and hippocampus) and cortical networks support curiosity-enhanced memory?

Our next set of analyses focused on understanding the subcortical-cortical interactions to elucidate if, and when, the hippocampus and SN/VTA communicate to higher-order brain networks to support curiosity-enhanced memory. We focused these connectivity analyses first on the whole-network level (Figure 4) and then parcellated the subnetworks into the key regions that make up each network (Figure 5; see also Methods for details).

**Figure 4.**
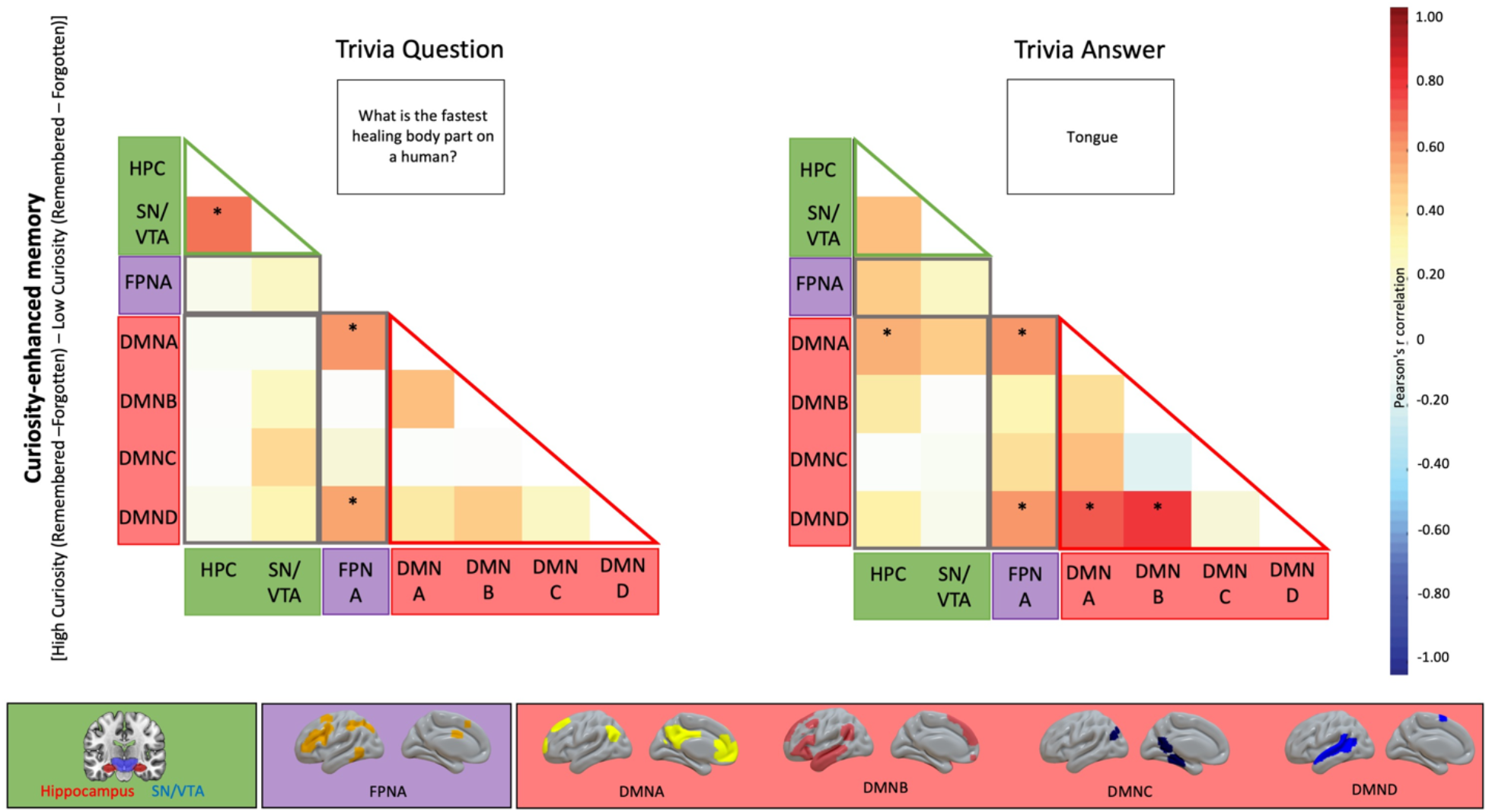
Subcortical connectivity (hippocampus - SN/VTA) correlated with curiosity-enhanced memory during the elicitation of curiosity, whereas between subcortical-cortical connectivity (hippocampus – DMNA) correlated with curiosity-enhanced memory during relief of curiosity. PPI analysis to investigate whether hippocampal-dopaminergic ROIs (HPC = hippocampus, SN/VTA = substantia nigra and ventral tegmental area (green)) showed increased functional correlations with the five cortical ROIs that showed a significant curiosity-enhanced memory effect (Figure 3; FPNA A (purple); DMNA; DMNB; DMNC and DMND (all DMN in red); highlighted in the bottom panel). All *p*-values were adjusted to control for false-discovery rate (FDR) using the Benjamini Hochberg correction in R. * indicates significant adjusted *p*-values < .05.

**Figure 5.**
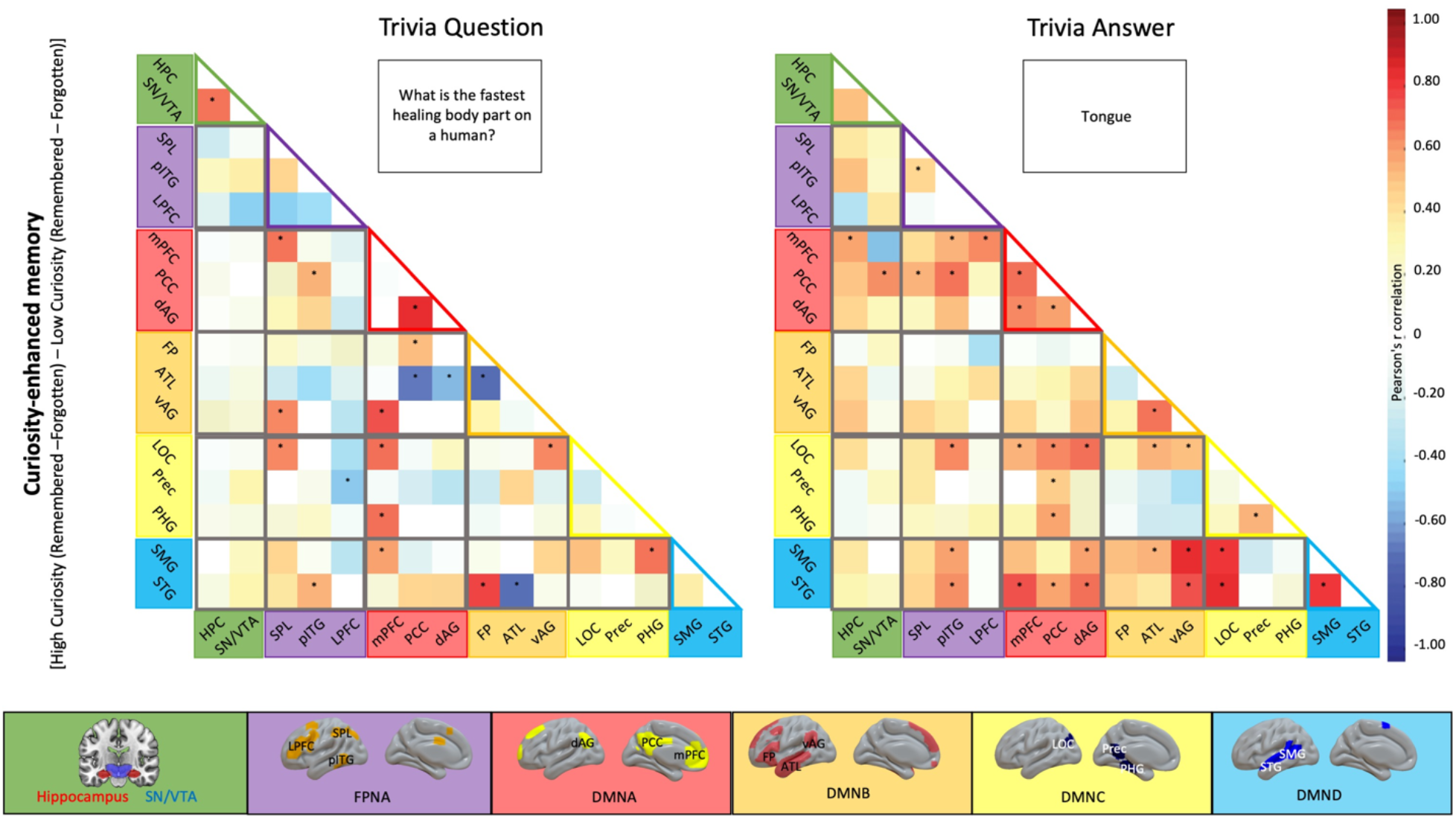
During the elicitation of curiosity, functional subcortical connectivity (hippocampus - SN/VTA) and within cortical networks (FPNA - DMN subnetworks) – but not subcortical-cortical coupling – correlated with curiosity-enhanced memory. However, during the relief of curiosity, coupling between subcortical regions and DMNA emerged in support of curiosity-enhanced memory (i.e., via SN/VTA-PCC and hippocampal-vmPFC coupling). PPI analysis to investigate whether hippocampal-dopaminergic regions (HPC = hippocampus, SN/VTA = substantia nigra and ventral tegmental area ROIs (green)) showed increased functional correlations with the five ROIs that showed a significant curiosity-enhanced memory effect (Figure 3; FPNA A; DMNA; DMNB; DMNC and DMND). For each of the five network ROIs we further parcellated these down into key regions (highlighted in the bottom panel). This resulted in the following ROIs for each subnetwork; FPNA in purple (SPL = superior parietal lobule, pITG = posterior inferior temporal gyrus, lPFC = lateral prefrontal cortex), DMNA in red (mPFC = medial prefrontal cortex, PCC = posterior cingulate cortex, dAG = dorsal angular gyrus), DMNB in orange (FP = frontal pole, ATL = lateral temporal lobe extending into anterior temporal lobe, vAG = ventral angular gyrus), DMNC in yellow (LOC = superior lateral occipital cortex, Prec = precuneous, PHG = parahippocampal gyrus) and DMND in blue (SMG = supramarginal gyrus, STG = superior temporal gyrus). The analysis was identical to that described in Figure 4 (see Methods for details). All *p*-values were adjusted to control for false-discovery rate (FDR) using the Benjamini Hochberg correction in R. * indicates significant adjusted *p*-values < .05.

#### During curiosity elicitation, only within subcortical-subcortical and within cortical-cortical functional connectivity (but not between subcortical-cortical connections) predict curiosity-enhanced memory

Consistent with previous findings (Gruber et al., 2014), we found that curiosity elicitation (i.e., during the presentation of the trivia question) elicited positive subcortical-subcortical functional connectivity. That is, we found that question-elicited functional connectivity between the hippocampus and SN/VTA ROI predicted curiosity-enhanced memory (Figure 4). Notably, curiosity-enhanced memory was not supported by any between subcortical-cortical interactions during the trivia question presentation in the network-wide (Figure 4A) and the parcellation analysis (Figure 5A). For the within cortical-cortical interactions during curiosity elicitation, the FPNA network ROI showed robust coupling with both DMNA and DMND in the network-wide analysis (Figure 4A) and also in the parcellation analysis (Figure 5A). We also found a non-significant positive trend between DMNA-DMNB in the network-wide analysis (Figure 4A). When we further investigated this finding in our parcellation analysis, we found that this trend was driven by a positive posterior cingulate cortex (PCC) – frontal pole (FP) coupling and a positive medial prefrontal cortex (mPFC) – ventral angular gyrus (vAG) coupling, but negative coupling between PCC and anterior temporal lobe (ATL), and negative coupling between ATL and dorsal angular gyrus (dAG) (Figure 5A). In addition, the parcellation analysis revealed that FPNA showed a more fine-grained connectivity with a portion of DMNC (i.e., lateral occipital cortex [LOC]) and DMN B (i.e., ventral angular gyrus [vAG]) that was not seen in the network-wide analysis. Finally, additional significant decoupling correlated with curiosity-enhanced memory: specifically, a portion of the DMNB (i.e., ATL) and DMND (j.e., superior temporal gyrus [STG]) (Figure 5A).

#### During curiosity relief, between subcortical-cortical connections (but not within subcortical-subcortical connections) predict curiosity-enhanced memory

During curiosity relief (i.e., the presentation of the trivia answer), no subcortical-subcortical (i.e., SN/VTA-hippocampus) connectivity correlated with curiosity-enhanced memory performance (Figure 4B, 5B). However, in our network-wide analysis the hippocampus ROI significantly coupled with the DMNA in support of curiosity-enhanced memory (Figure 4B). When we followed this up with our parcellation analysis during curiosity relief, it revealed that the hippocampus significantly coupled with the mPFC (Figure 5B). Furthermore, while SN/VTA did not couple with an entire network, the parcellation analysis revealed that the SN/VTA ROI did significantly couple with the PCC of DMNA in support of curiosity-enhanced memory (Figure 5B). Furthermore, there was substantial cortical-cortical connectivity in support of curiosity-enhanced memory (Figure 5B). Comparable to the trivia question presentation, network-wide analysis identified that FPNA showed significant coupling to DMNA and DMND during curiosity relief (Figure 4B). The further parcellation analysis revealed that this was driven by two key regions of FPNA (i.e., posterior inferior temporal cortex [pITC] and superior parietal lobule [SPL]), coupling with regions of DMNA (i.e., PCC and dAG) and DMND (i.e., SMG and STG) (Figure 5B). Finally, during trivia question presentation, extensive significant connectivity within and across DMN subnetworks was seen to support curiosity-enhanced memory (Figure 5B). We did not find any significant decoupling in support of curiosity-enhanced memory during answer presentation.

## Discussion

The current fMRI study used a trivia paradigm to examine the role of the FPN and DMN in supporting curiosity-enhanced memory. We identified that specific recruitment of FPNA and widespread recruitment of all DMN subnetworks support curiosity-enhanced memory at both the elicitation (trivia question presentation) and relief of curiosity (trivia answer presentation). Furthermore, we investigated functional connectivity between subcortical regions previously implicated in curiosity-enhanced memory (SN/VTA and hippocampus) and the FPN and DMN subnetworks. These results indicate a timepoint specific effect on when subcortical regions (SN/VTA and hippocampus) couple with themselves and to cortical regions. During curiosity elicitation, within subcortical-subcortical (SN/VTA–hippocampus) and within cortical-cortical functional connectivity (FPNA–DMNA; FPNA–DMND), but not between subcortical-cortical connections, predict curiosity-enhanced memory. In contrast, during curiosity relief, between subcortical-cortical connections (via SN/VTA-PCC and hippocampal-vmPFC coupling) and within cortical-cortical functional connectivity (FPNA–DMNA; FPNA-DMND; DMND–DMNA and DMNB), but not within subcortical-subcortical connections, predict curiosity-enhanced memory.

### FPN supports curiosity-enhanced memory independently of subcortical connectivity, but functionally couples with DMN subnetworks in support of curiosity-enhanced memory

We found increased BOLD response activation for high- compared to low-curiosity conditions in one of the two frontoparietal networks – FPNA (but not FPNB). This network has been demonstrated to be crucially involved in semantic control (Davey et al. 2015; Whitney et al. 2011), which enables us to flexibly access and manipulate meaningful information to focus on the aspects of a concept that are relevant to a particular context or task (Jackson, 2021). Consistent with this idea, we found curiosity-related activation in FPNA during curiosity elicitation (potentially suggesting the flexible access to semantic information) and during curiosity relief (suggesting the manipulation and comparison of new information with stored knowledge). Furthermore, increased activation in FPNA, but not FPNB, was associated with curiosity-enhanced memory. Our findings therefore extend recent findings by Duan and colleagues (2021), who showed activity in a set of regions overlapping with the frontoparietal network that supported curiosity-enhanced memory, by further pinpointing that this effect seems to be specific to areas supporting semantic control.

In addition, our functional connectivity analyses revealed that FPNA did not show coupling with SN/VTA or hippocampal regions to support curiosity-enhanced memory at either elicitation or relief. However, robust coupling between FPNA and both DMNA and DMND was seen at both timepoints. The coupling between control and DMN regions is consistent with findings that functional connectivity between FPN and DMN contributes to a ‘curiosity network’, with PFC (core region of FPNA) and angular gyrus (core region of DMNA) making the greatest contribution to prediction power of trait curiosity (Li et al., 2019). A recent study also highlighted that FPN neurons anticipate information about rewards (Jezzini et al., 2021), while DMN regions play a role during encoding and retrieval (Maillet & Rajah, 2014) to enable integration of information into contextual schemas over time. We therefore speculate that the increased connection between FPN and DMN might reflect the increased synergy of cognitive control, information retrieval, and integration that leads to enhanced memory for information associated with high curiosity.

#### DMN subnetworks differentially support general curiosity and curiosity-enhanced memory

Participants recruited different DMN subnetworks associated with curiosity compared to curiosity-enhanced memory. DMNA did not dissociate between high- and low-curiosity conditions but was modulated by curiosity-enhanced memory. DMNA corresponds to the core default mode network and is most heavily associated with introspective states such as self- orientated thought and mind-wandering (Raichle et al., 2001; Stawarczyk et al., 2011). Recent perspectives have also suggested this network is an active and dynamic ‘sense-making’ network that integrates incoming extrinsic information with prior intrinsic information to form rich, context-dependent models of the world as it unfolds over time (Stawarczyk et al., 2021; Yeshurun et al., 2021). Furthermore, the human lesion literature has reported that robust memory impairment can be observed following mPFC damage (a key region of DMNA), suggesting that the mPFC has a role in initiating and coordinating cognitive processes for episodic simulation (McCormick et al., 2017). This perspective suggests that DMNA recruitment enables participants to more successfully accumulate and integrate new trivia information into their prior schemas for later remembered high-curiosity information.

A more domain general role was seen for DMNB, as this network supported both states of curiosity and curiosity-enhanced memory in our BOLD activation analysis. DMNB aligns with the semantic network (Jackson et al., 2019; Lambon Ralph et al., 2017), which has been shown to support the combination of concepts into meaningful and more complex representations (Pylkkänen, 2019). A recent study showed conceptual information related to the current goal dominates the multivariate response within DMNB, which supports flexible retrieval by modulating its response to suit the task demands (Wang et al., 2021). As curiosity may be induced through individuals’ awareness of the discrepancy between their current informational and goal uncertainty states (Gottlieb et al., 2013), curiosity can be seen as a motivational state that seeks information to fill this knowledge gap. Taken together, this suggests that DMNB activation might be coding the intrinsic goal (i.e., the goal of curiosity is to seek information to fill knowledge gap) and combines the trivia information into meaningful representations, thereby supporting curiosity-enhanced memory.

Interestingly, a significant functional decoupling between DMNB (in particular, anterior temporal lobe) and both DMNA (dorsal angular gyrus and posterior cingulate cortex) and DMND (superior temporal gyrus) correlated with curiosity-enhanced memory during the elicitation of curiosity. The shared pattern of decoupling between DMNA and DMNB has been previously associated with better verbal semantic performance (Vatansever et al., 2017). Segregation of these regions during the elicitation of curiosity may therefore be crucial for the capacity to engage successfully in upcoming externally-presented high-curiosity information while simultaneously accessing the semantic store to determine the exact knowledge gap.

Finally, there was also substantial functional interconnection across DMN subnetworks during trivia question presentation, seen to support curiosity-enhanced memory. It is argued these networks work in tandem to support the retrieval of content during naturalistic events. For example, it has previously been shown that both DMNB show intercommunication with language networks (Gordon et al., 2020), while DMNA may integrate and coordinate information across the whole DMN (Barnett et al., 2021), bolstering the idea that interconnection of DMN subnetworks facilitates higher-order integration and coordination of memory-guided activity.

#### During curiosity relief, hippocampus and SN/VTA couple specifically with DMNA in support of curiosity-enhanced memory

Broadly consistent with previous findings on curiosity-enhanced memory (Gruber et al., 2014; Poh et al., 2021), we found that during curiosity elicitation – but not during curiosity relief – positive subcortical-subcortical functional connectivity between the hippocampus and SN/VTA region predicted curiosity-enhanced memory. Our findings draw parallels with those from the reward-motivated memory literature that have shown that increased functional connectivity between the SN/VTA and hippocampus supports reward-related memory benefits (Adcock et al., 2006, Gruber et al., 2016; Murty & Adcock, 2014). Perhaps surprisingly, we did not find that any subcortical-cortical interactions during curiosity elicitation supported curiosity-enhanced memory. Based on the Prediction, Appraisal, Curiosity, and Exploration (PACE) framework (Gruber & Ranganath, 2019) and other findings on anticipatory reward states (Marvin et al., 2020; Murty & Adcock, 2014; Murty, Ballard & Adcock, 2017), we would have expected that lateral PFC (a key region of the FPN) and perhaps more widely the FPN would couple with the SN/VTA in support of curiosity-enhanced memory. However, in the present study, we passively presented participants with high- and low-curiosity trivia questions without the need to make any decisions regarding the information they were encoding. Future research on curiosity-enhanced memory would need to use paradigms that are sensitive to whether/how appraisal-based lateral prefrontal input couples with the SN/VTA or the hippocampus to support curiosity-enhanced memory.

Critically, specifically during curiosity relief, we found that the hippocampus significantly coupled with the DMNA – particularly with the mPFC – in support of curiosity-enhanced memory. This finding is consistent with previous findings on how instantiation and reinstatement of schemas are mediated by the interaction among regions within the DMN and hippocampus (Gilboa & Marlatte, 2017; Ranganath & Ritchey, 2012; Yeshurun, Nguyegn & Hasson, 2021). Consistent with prior evidence suggesting that BOLD signal fluctuations in the DMN are tighly functionally connected to dopaminergic midbrain regions (e.g., SN/VTA) (Bär et al., 2016), we found that SN/VTA significantly coupled with PCC of DMNA in support of curiosity-enhanced memory. Taken together, this provides a plausible neuromodulatory mechanism through which hippocampal-dopaminergic dynamics elicit curiosity and subsequently communicate to higher-order brain regions within DMNA to facilitate curiosity-enhanced memory integration.

## Conclusion

In summary, our findings demonstrate how two key cortical networks related to semantic control and memory integration (i.e., the FPN and DMN, respectively) support curiosity and curiosity-enhanced memory via FPN-DMN interactions and DMN subnetwork interactions. Here, we show how the dopaminergic system together with the hippocampus interact specifically with subnetwork DMNA potentially reflecting how subcortical regions support the enhancement of memory intergration of semantic information associated with high curiosity. Our findings suggest that future research on curiosity would need to better understand how cortical networks indepdently and jointly with subcortical systems interact to faciliate memory formation for high-curiosity information.

## Acknowledgments

This study was supported by a Wellcome Trust and Royal Society Sir Henry Dale Fellowship (211201/Z/18/Z) awarded to M.J.G.

